# A novel micro-ECoG recording method for recording multisensory neural activity from the parietal to temporal cortices in mice

**DOI:** 10.1101/2022.10.01.510247

**Authors:** Susumu Setogawa, Ryota Kanda, Shuto Tada, Takuya Hikima, Yoshito Saitoh, Mikiko Ishikawa, Satoshi Nakada, Fumiko Seki, Keigo Hikishima, Hideyuki Matsumoto, Kenji Mizuseki, Osamu Fukayama, Makoto Osanai, Hiroto Sekiguchi, Noriaki Ohkawa

## Abstract

Characterization of inter-regional interactions in brain is essential for understanding the mechanism relevant to normal brain function and neurological disease. The recently developed flexible micro (μ)-electrocorticography (μECoG) device is one prominent method used to examine large-scale cortical activity across multiple regions. The sheet-shaped μECoG electrodes arrays can be placed on a relatively wide area of cortical surface beneath the skull by inserting the device into the space between skull and brain. Although rats and mice are useful tools for neuroscience, current μECoG recording methods in these animals are limited to the parietal region of cerebral cortex. Recording cortical activity from the temporal region of cortex in mice has proven difficult because of surgical barriers created by the skull and surrounding temporalis muscle anatomy. Here, we developed a sheet-shaped 64-channel μECoG device that allows access to the mouse temporal cortex, and we determined the factor determining the appropriate bending stiffness for the μECoG electrode array. We also established a surgical technique to implant the electrode arrays into the epidural space over a wide area of cerebral cortex covering from the barrel field to olfactory (piriform) cortex, which is the deepest region of the cerebral cortex. Using histology and computed tomography (CT) images, we confirmed that the tip of the μECoG device reached to the most ventral part of cerebral cortex without causing noticeable damage to the brain surface. Moreover, the device simultaneously recorded somatosensory and odor stimulus-evoked neural activity from dorsal and ventral parts of cerebral cortex in awake and anesthetized mice. These data indicate that our μECoG device and surgical techniques enable the recording of large-scale cortical activity from the parietal to temporal cortex in mice, including somatosensory and olfactory cortices. This system will provide more opportunities for the investigation of physiological functions from wider areas of the mouse cerebral cortex than those currently available with existing ECoG techniques.

## Introduction

Neural networks in the cerebral cortex are composed of functionally different regions which process information from distinct modalities to drive cognitive functions including attention, learning, and memory. Altered network activity from multiple cortical regions is linked to neurological disorders, such as seizure, dementia, and schizophrenia [1–3]. Therefore, characterization of inter-cortical interactions is essential for understanding the mechanisms underlying normal cognitive functions and neuropathology.

Methods in neuroscience, such as large-scale calcium imaging, electroencephalography (EEG), and electrocorticography (ECoG), enable the monitoring of neural activity from a wide area of cerebral cortex, providing insight into inter-regional cortical network interactions [4–6]. These techniques, combined with rodent experiments and a variety of genetic tools, have contributed to advances in neuroscience[7]. However, the measurement area of current large-scale calcium imaging and electrophysiological techniques focusing on cerebral cortex has been mostly limited to parietal regions, including sensorimotor, visual, auditory, parietal, and retrosplenial cortices [5, 8, 9]. A major reason for the limitation in measurement area is the difficulty in the associated operation, since the temporal region of the cortex is covered by skull surrounded with temporalis muscles [10]. Moreover, the most common methods that use head fixation in rats and mice, such as the ear bars used in craniotomy and the head-fixation plate[11], interferes with microscopy and electrode implantation for large-scale recordings of the temporal cortex. Given that the temporal cortex in rodents, including the piriform, insular, and entorhinal cortex, is involved in various types of cognitive processes [12–14], establishing large-scale recording techniques covering the parietal to temporal parts of cortices would increase understanding of network dynamics underlying normal cognitive functions and their disruption in diseases.

The μECoG device with high-density multi-channel electrodes and fabricated by semiconductor technology enables the simultaneous recording of local field potentials (LFPs) from multiple regions in cerebral cortex with high spatiotemporal resolution [15–17]. The sheet-shaped μECoG device which has been developed using parylene C to reduce sheet thickness, has improved electrode adhesiveness to the curved surface of the brain[6]. Thinning of the sheet-shaped electrodes reduces stiffness and enhances electrode contact to the subject [18]. It has been shown that a relatively thick sheet of μECoG electrode array can be implanted into the cerebral cortex through a small slit on the parietal bone [17, 18]. The electrode arrays make good contact with the brain surface due to compression pressure between the brain and skull, despite adopting the relatively thick sheet in the electrode array. Previous experiments have improved μECoG techniques in order to mitigate anatomical and environmental factors that hinder electrophysiology recordings from cortical regions. Whereas μECoG techniques have the potential to provide a more large-scale neural recording system, a method that covers the parietal and temporal cortex in rodent models has not been developed.

The goal of this study is to develop a sheet-shaped μECoG device that is implantable into the epidural space, and to establish the optimal surgical procedures to record from the multisensory regions in temporal cortex. From calculations on the bending stiffness of the μECoG sheet, we showed that the thickness, width, and channel number of metal wiring are key factors for developing μECoG parylene-based electrode arrays. These arrays can be inserted into the space between the skull and brain for optimal large-scale neural recording. After surgical insertion of the sheet-shaped μECoG device, we used histological methods and CT images to confirm that the position of the sheet tip reached to the most ventral part of cerebral cortex without structural damage to the brain surface. This μECoG device allowed successive LFP recordings along the dorsoventral arc of cerebral cortex in an awake mouse. In addition, we simultaneously recorded neural responses corresponding to somatosensory and odor stimuli from different electrodes placed on the parietal and temporal cortex in anesthetized mice. These results show that the sheet-shaped μECoG permits acquisition of large-scale LFP recordings from the parietal to temporal cortex in mice, including somatosensory and olfactory areas.

## Materials and methods

### Device fabrication

The neural electrode array was fabricated on biocompatible parylene film using a standard semiconductor process (Fig. 1a). First, a thick parylene C film was deposited as a base layer on a Si substrate using a parylene coater (PDS2010, Specialty Coating Systems Inc.). Subsequently, a Ti/ Au (50 nm/200 nm) metal wiring layer, including metal neural electrode array, was deposited on parylene film using an electron beam evaporator (ED-1600, SANVAC Co., Ltd.). The array consists of 64 Ti/Au metal electrodes with an area of 80 × 80 μm^2^ in a 32 (300 μm pitch) × 2 (400 or 800 μm pitch) configuration. Limited channel crosstalk from the electrode arrangement was confirmed using an equivalent circuit model (Additional File: Additional Method, Fig. S1). The interconnected metal wiring layer was covered with 1 μm thick parylene film. A focal area of parylene was removed by CF_4_ and O_2_ plasma gas etching to form apertures on the Ti/Au electrode (RIE-200NL, Samco Inc.). Finally, the electrode array was mounted on a printed circuit board (PCB) with a design of 15.5 mm × 14.3 mm using flip chip bonder (M90, HiSOL, Inc.). The cross-section of the electrode array, which is placed on cerebral cortex, is shown in Fig. 1b.

**Fig. 1.**
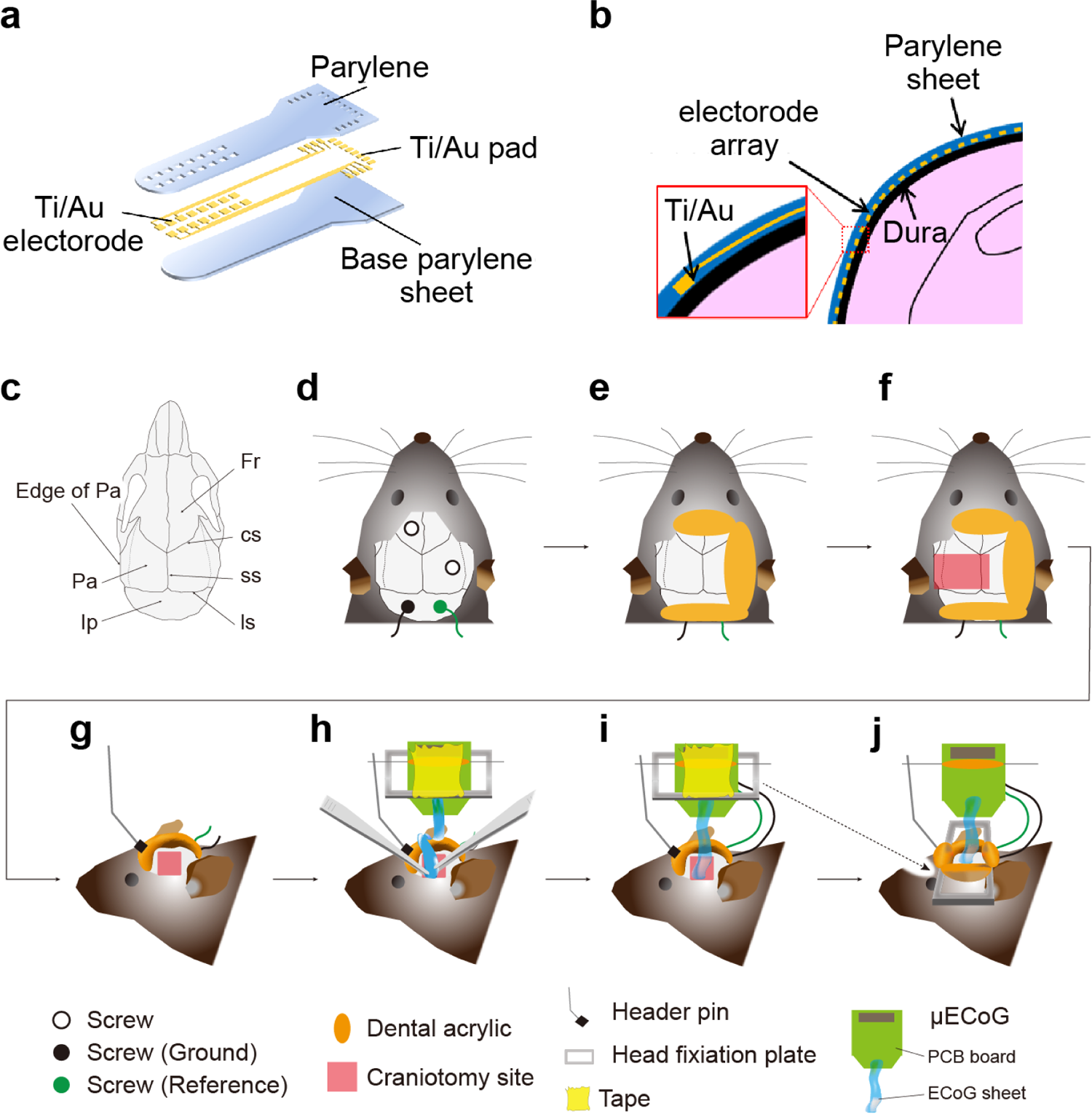
Surgical procedures for the placement of the sheet-shaped μECoG device. (a) A schematic of the device showing its various components. (b) Schematic representation of a cross-section of the neural electrode array device placed on the cerebral cortex. (c) The cranial sutures in the dorsal view, Fr, Frontal bone; IP, Interparietal bone; Pa, parietal bone; cs, coronal suture; ss, sagittal suture; ls, lambdoid suture; edge of Pa, the edge of parietal bone connecting the temporalis muscle. (d-j) Schematic representations of the surgical procedures for inserting the μECoG electrode array into the space between the brain and skull.

### Animals

Native male mice C57BL6/J (Japan SLC, Inc., Shizuoka, Japan) were used at 3– 6 months of age. All mice were maintained on a 12 h light/dark cycle (lights on 7:00 am– 7:00 pm) at 24 ± 3 °C and 55 ± 5% humidity, had ad libitum access to food and water, and were housed in a cage with littermates until surgery.

### Surgical implantation of the flexible **μ**ECoG sheet

Male mice (n = 5, 31.1 ± 2.1 g) were implanted with μECoG arrays in the left cranial hemisphere. During surgery, mice were anesthetized with 0.75 mg/kg medetomidine hydrochloride, 4.0 mg/kg midazolam, and 5.0 mg/kg butorphanol tartrate for recordings under awake conditions, or urethane (1.2 g/kg) for recordings under anesthetized conditions. Mice were placed into a stereotaxic frame (Narishige, Tokyo, Japan) and on a heating pad to maintain normal body temperature (37 °C; MK-900, Muromachi Kikai, or ATC-TY, Unique-Medical Inc., Tokyo, Japan) during surgery. An incision was made in the scalp along the midline of the brain. After exposing the skull, a tendon of temporalis muscle connecting to the edge of parietal bone was cut to secure a space to perform implantation of the μECoG array (Fig. 1c-j). Four stainless steel anchor screws (AN-3, Eicom, Kyoto, Japan) were attached to the skull for mechanical support; one screw was placed on the frontal bone and one on the parietal bone, while the remaining screws, which served as ground and reference, were inserted into the interparietal bone above cerebellum (Fig. 1d). Before the craniotomy, dental acrylic (UNIFAST III, GC Corporation, Tokyo, Japan) was placed on the edge of the exposed skull to affix screws and to protect ground and reference wires (Fig. 1e). The skull was drilled into a rectangular shape from bregma to lambda on the anteroposterior axis and from the right parietal bone to the left squamosal bone on the mediolateral axis (Fig. 1f, approx. 4.5 mm × 6 mm (anteroposterior axis × mediolateral axis)). The surgical procedures ensured preservation of the dura. The head-fixation plate was taped onto the PCB in order to affix it on the head after implantation of the μECoG sheet. A male header pin (TSW-105-20-T-S, DegiKey, USA) was attached to the frontal bone for a device holder, and the μECoG was soldered to secure the PCB onto the skull (Fig. 1g). The μECoG array was carefully inserted into the space between the brain and squamosal bone (Fig. 1h). Note the flexible ECoG sheet was placed on the dura mater. After implantation, the craniotomy site was covered by the parietal bone that was removed during craniotomy, and was embedded using dental acrylic (Fig. 1i). A head-fixation plate, temporarily supported with tape on the PCB, was attached on the head (Fig. 1j, dashed line). After the surgery, mice were placed into a home cage and on a heating pad to maintain body temperature until recordings. Recordings from awake mice were conducted at least 3 hours after the completion of surgery. Note that the most important point for successful implantation is to avoid damage to the dura matter. This was achieved by carefully manipulating the drill, tweezer, and device during surgical procedures (Fig. 1f-h).

### Behavioral task and recordings

Mice were head-fixed on the apparatus and placed in a sound attenuated chamber for the multisensory presentation task. An electrical fan provided background sound (50 dB) and ventilation. White LED was placed in front of the right eye, and odor and air tubes were placed in front of the nose and whiskers, respectively. The controller for multisensory presentations was developed using custom programs (LabView, National Instruments, TX). The program pseudorandomly controlled the visual (Frequency, 10 Hz), somatosensory (Flow rate, 2.5 L/min), and odor (Flow rate, 1 L/min, 1% benzaldehyde diluted in mineral oil) stimuli for 3 sec with intertrial intervals (∼10–20 sec) that lasted for 100 or 150 trials in the awake or in the anesthetized experiment, respectively (50 trials each stimulus).

LFP and event signals were recorded at a sampling rate of 20 kHz using the Open Ephys recording system (www.open-ephys.org). Data were downsampled to 1,250 Hz for analysis. Neural signals were amplified on the 64-Channel head stage (C3325, Intan Technologies, USA). To connect the μECoG device and the head stage, a custom-made adapter corresponding to both the flexible flat cable connector (39FVXS-RSM1-GAN-TF, J. S. T. MFG. Co., Ltd, Japan) on the device and the nano strip connectors on the head stage (C3315, Omnetics, USA) was used. Data were analyzed using MATLAB software (MathWorks, MI, USA). The stimulus-evoked LFP was obtained by averaging the LFPs against stimulus onset across trials. To identify sensory stimulus-evoked LFPs on active channels, we compared the peak LFP amplitudes between the before (−600 to 0 ms) and after (0 to 600 ms) stimulus for each electrode. To compare the LFP amplitude between the different electrodes or stimuli, we also used the peak LFP amplitudes for 0.6 sec after stimulus onset. To avoid issues caused by the current transmitting far from the μECoG device, we obtained the profiles of LFPs (LFP profile) by calculating the second spatial derivative of the LFPs [19, 20]. The second spatial derivative is approximated by 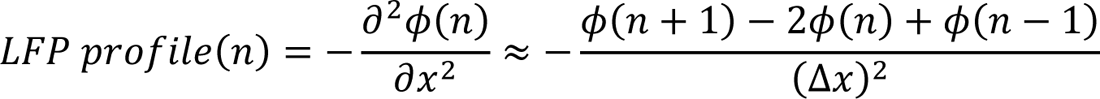 where ϕ is the LFP, Δx is the distance between adjacent electrodes of the μECoG device, and n is the electrode number. The electrode position with a larger LFP profile amplitude indicates the location with a higher current flow than the surrounding regions, indicating that the neural activity near the electrode with a higher LFP profile is high. The LFP profile was calculated using the MATLAB script written by Timothy Olsen (2022) (https://www.mathworks.com/matlabcentral/fileexchange/69399-current-source-density-csd).

### Histological analysis

After the μECoG implantation and recordings, mice were deeply anesthetized with 5% isoflurane and an overdose of pentobarbital solution, and then transcardially perfused with 0.9% saline. To check the position of the μECoG sheet, the head was cut, and the muscles surrounding the skull were removed using a bone rongeur. The skull was quickly covered in crushed dry ice for flash-freezing. The frozen skull was stored at −30 ℃ until use. To prepare sections including brain, skull, and the flexible μECoG sheet, we used a polyvinylidene chloride film and a modification of a previously described method [21]. The skulls were embedded in OCT compound at −20°C for at least 20 min before slicing in the cryostat. After reaching to the sampling coordinates, the coronal surface of the brain was covered by transfer film (Cryofilm type ⅡC(9), SECTION-LAB, Hiroshima, Japan) to avoid detaching the skull, brain, and μECoG sheet from samples. Samples were cut at 30 μm. Serial sections were placed on glass slides (Matsunami, Osaka, Japan) and stained with 4’,6-diamidino-2-phenylindole (DAPI; D9542, Sigma-Aldrich, St. Louis, MO) for 5 min. Sections were rinsed by phosphate buffered saline for 3 min and allowed to dry for 20 min. The films containing the sections were flipped onto a drop of embedding agent (ProLong antifade reagents, Thermo Fisher Scientific, USA) on a new glass slide and gently pushed with a finger to adhere them to the substrate. The redundant embedding agent was removed by filter paper, and the perimeter of the film was covered with manicure to prevent tissue drying. Sections were scanned using a BZ-X700 (Keyence, Osaka, Japan) with 4 × objective lens. The contrast and color balance of images scanned under bright field and UV light conditions were adjusted using Photoshop (CC, Adobe Inc., USA). The images were superimposed for analysis.

### CT imaging

To visualize the position of the μECoG electrode array without removing the skull, we used CT scans (Cosumo Scan Fx, Rigaku Corp., Tokyo Japan). After recordings, mice were perfused with 0.9% saline as described under histological analysis. The PCB of the μECoG and header pin were removed from the head of mice, and the remaining ECoG sheet was bonded to the skull to avoid displacement of the electrode position. The CT data were obtained using the following parameters; tube voltage, 90 kV; tube current, 88 μA; exposure time, 2 min; and resolution 20 μm isotropic.

### Statistics

Statistical analysis was performed using MATLAB software. Comparison of data between two groups was performed using the Student’s *t*-test (paired), the Wilcoxon signed rank test (paired), and the Wilcoxon rank-sum test (unpaired). Comparison of data between three groups were constructed using a one-way repeated measures ANOVA or the Friedman test (without assuming normal distribution and homogeneity of variances). The level of statistical significance for variables was set at *p < 0.05, **p < 0.01, ***p < 0.001, †p < 0.05, †††p < 0.001. All statical tests were two-sided.

## Results

### Device fabrication

The thickness of the film is an important factor of stiffness for proper device insertion into the epidural space between the temporal cortex and skull. Devices with different thicknesses were fabricated to understand the effect of parylene thickness on the self-supporting characteristics. The parylene thicknesses *T*_p_ of the devices prepared in this study were 7, 13, and 23 μm. Fig. 2a shows photographs of the neural electrode array device with different thicknesses, including optical micrographs of the electrode array. The thin device was curled up owing to strain caused by the thermal expansion coefficient between metal and parylene. As increased thickness of the film results in increased bending stiffness, the thick device was not as curled, but resulted in more robust self-supporting characteristics relative to the thin device. The bending stiffness *S* of the parylene film without metal can be expressed as 

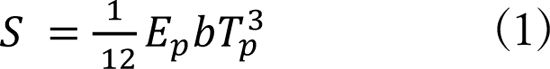

where *E*_p_ and *b* are the Young’s modulus (*E*_p_ = 2.78 GPa) and the width (*b* = 3.1 mm) of parylene film, respectively. It can be seen that *S* increases rapidly with increasing the *T*_p_. The actual device contains 64 channel Ti/Au metal wiring layer with a high Young’s modulus, which affects *S*. The structured-corrected *S* was theoretically calculated based on a previous report [22]. Fig. 2b shows the calculated bending stiffness of the device with and without the metal wiring layer as a function of the parylene thickness. The *S* of the device with the metal layer is ∼1.3–1.4 times higher than that of the device without the metal layer. To determine the optimal device design, the thickness and width for parylene, and also the thickness, width, and number of metal wire channels were key factors. Fig. 2c shows the frequency dependence of impedance from a representative neural electrode. Impedances at 1 kHz were approximately 0.5 MΩ. The distribution of channel impedance from the electrode used in the recording experiment in an awake mouse (Fig. 4) is shown in Fig. 2d (0.46 ± 0.07 MΩ). All the devices used (Fig. 5) showed similar impedance characteristics.

**Fig. 2.**
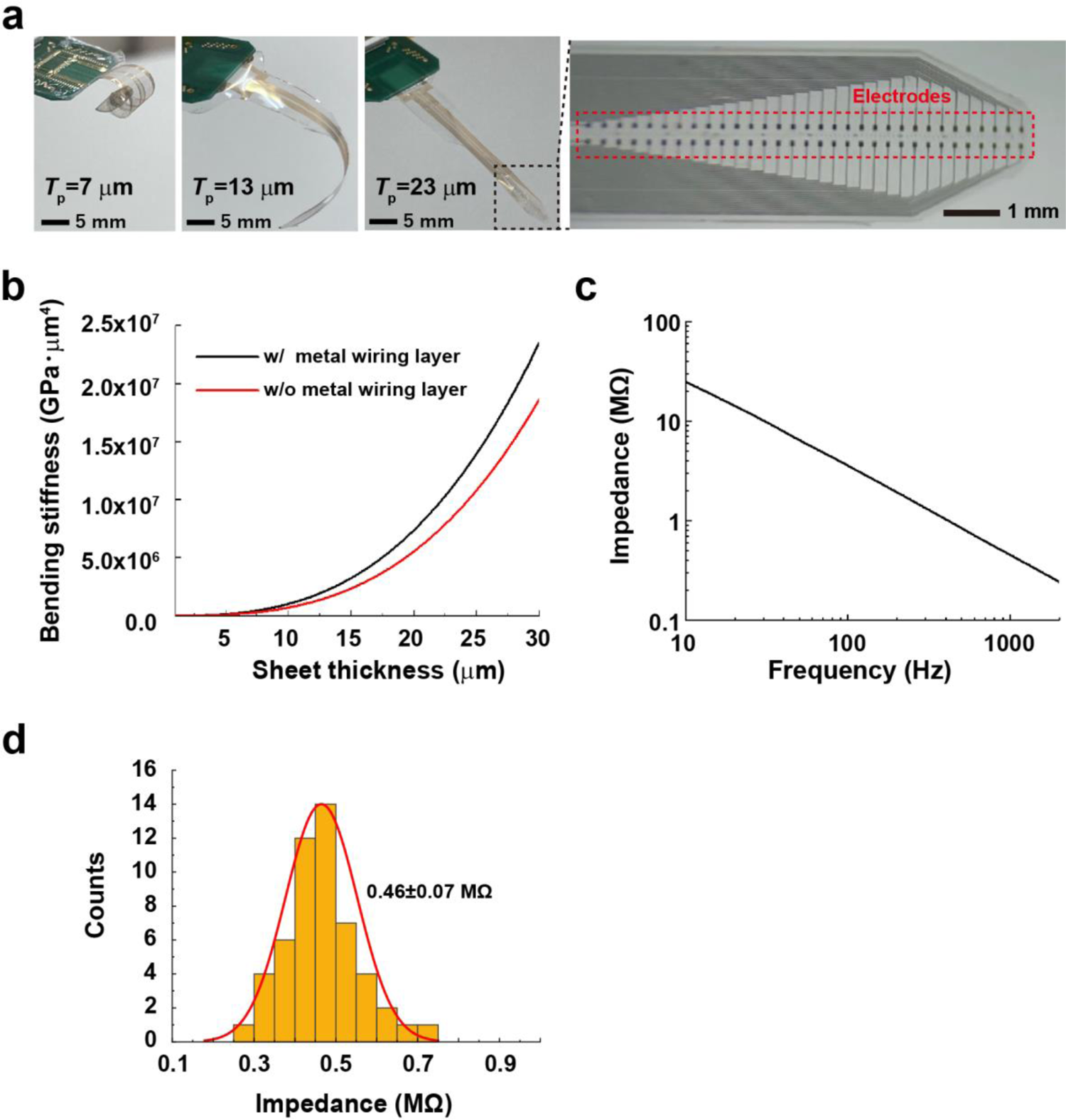
Bending stiffness of the sheet-shaped μECoG device. (a) Photographs of a μECoG sheet with different parylene thicknesses of 7, 13, 23 μm and an optical microscope image of a sheet with arranged neural electrodes (Ti/Au). While the 7 μm thick sheet strongly curls, the 23 μm thick sheet has high self-supporting characteristics. (b) Calculated bending stiffness of the device with and without the metal wiring layer as a function of the parylene thickness. (c) Frequency dependence of impedance of the representative neural electrode. The impedance at 1 kHz was approximately 0.5 MΩ. (d) The distribution of impedances of μECoG electrodes (52 / 64 channels). The remaining channels were removed due to high impedance (> 5 MΩ).

**Fig. 3.**
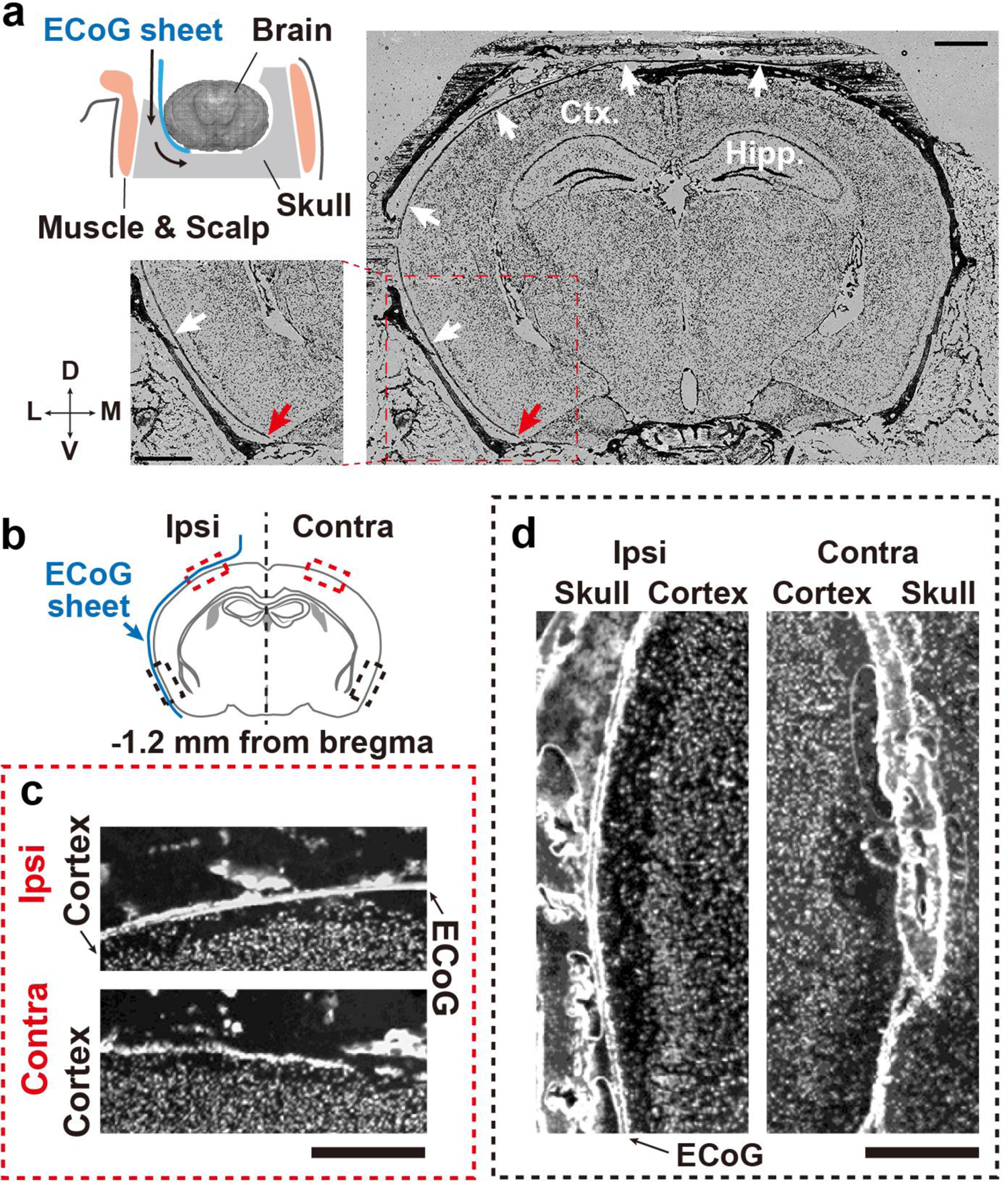
Histological validation of the μECoG sheet position. (a) A schematic of the position of the μECoG sheet placed on the brain surface (upper left). Superimposed microscopic images of bright field and DAPI staining (right and lower left). Black arrows indicate the μECoG sheet on the brain surface. The insets show enlargements of the dashed rectangle. Red arrows indicate the tip of the μECoG sheet. Scale bars: 1 mm. Ctx, cortex; Hipp, hippocampus; D, dorsal; M, medial; V, ventral; L, lateral. (b-d) Comparison of cortical structure between the μECoG sheet-inserted hemisphere (ipsi-lateral side) and opposite hemisphere (contralateral side). Ipsi, ipsilateral; Contra, contralateral. (b) Schematic drawing of a coronal view of the mouse brain; black and red dashed rectangles indicate the parietal (c) and temporal cortex (d), respectively. Scale bars: 500 μm.

**Fig. 4.**
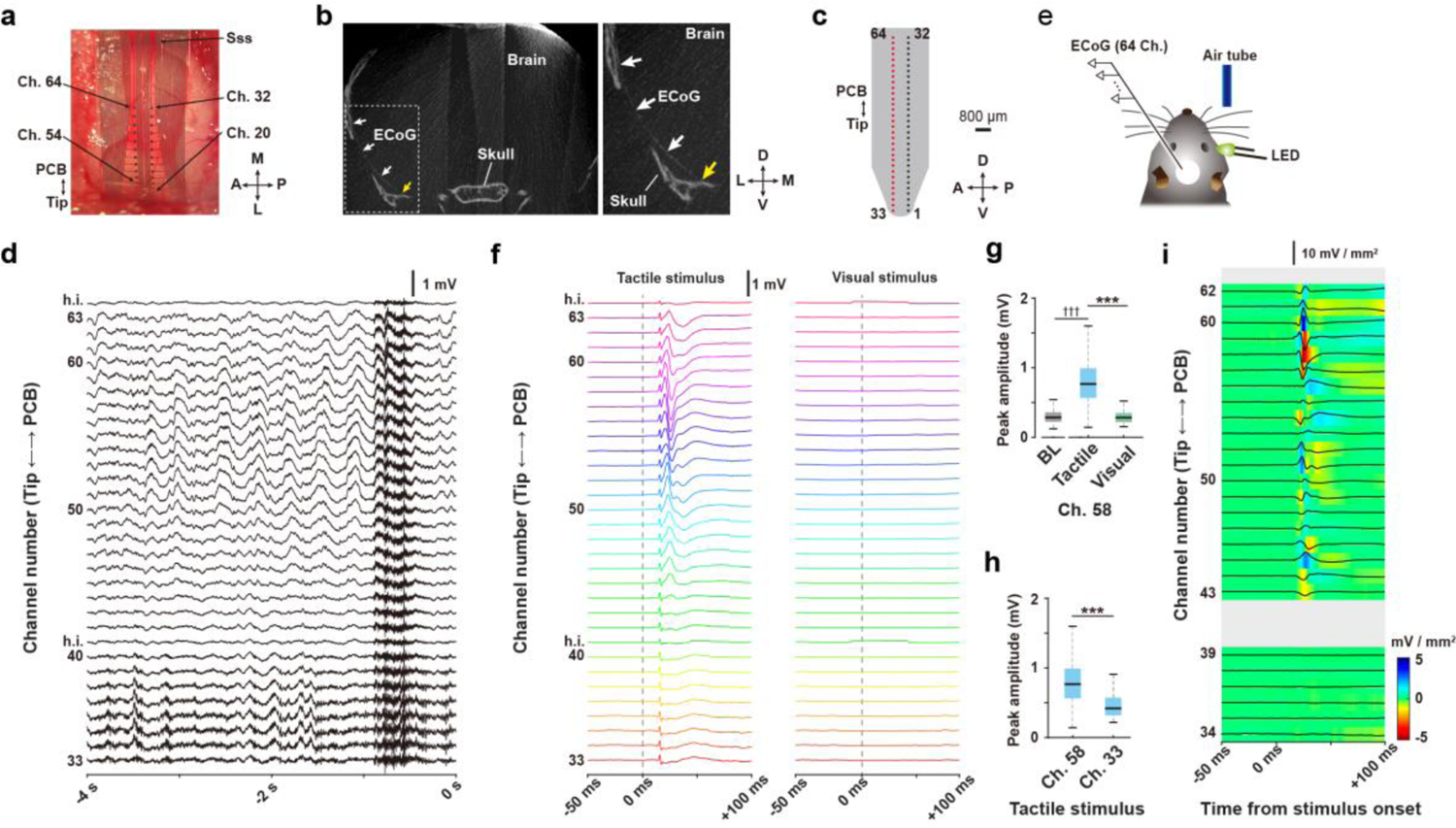
LFP Recordings by the μECoG sheet electrode in an awake mouse. (a) Dorsal view of the μECoG electrode array placed on the parietal cortex. M, medial; L, lateral; A, anterior; P, posterior; Ch., channel of electrode; Sss, superior sagittal sinus. (b) The position of the μECoG sheet placed on the brain surface as revealed by CT scanning. White arrows indicate the μECoG sheet placed on space between the brain and skull. The enlarged image (right) shows the tip of μECoG sheet in the brain (yellow arrow). D, dorsal; V, ventral; M, medial; L, lateral. (c) The 64-channel μECoG electrode array. Numbers are the channel IDs of the electrode. Red rectangles indicate the thirty-two successive electrodes that recorded representative LFPs along the dorsoventral axis of the cortex. D, dorsal; V, ventral; A, anterior; P, posterior. (d) Representative LFPs before the sensory presentation task. Numbers indicate channels of the electrode corresponding to (c) h.i., channel of high impedance (> 5 MΩ). (e) Schematic drawing of the sensory presentation task and recordings in a head-fixed mouse. (f) Average sensory-evoked LFPs. (g) Averaged peak LFP amplitudes in the same channels elicited by different stimuli. BL, baseline. (h) Averaged peak LFP amplitudes in different channels elicited by the same stimulus. (i) The current source density map calculated from the averaged tactile-evoked LFPs in (f). The central lines and edges of the boxes indicates the median and quartiles, respectively, and whiskers extend to the maximum and minimum. \*\*\**P* < 0.001. ††† < 0.001 vs. baseline amplitude.

**Fig. 5.**
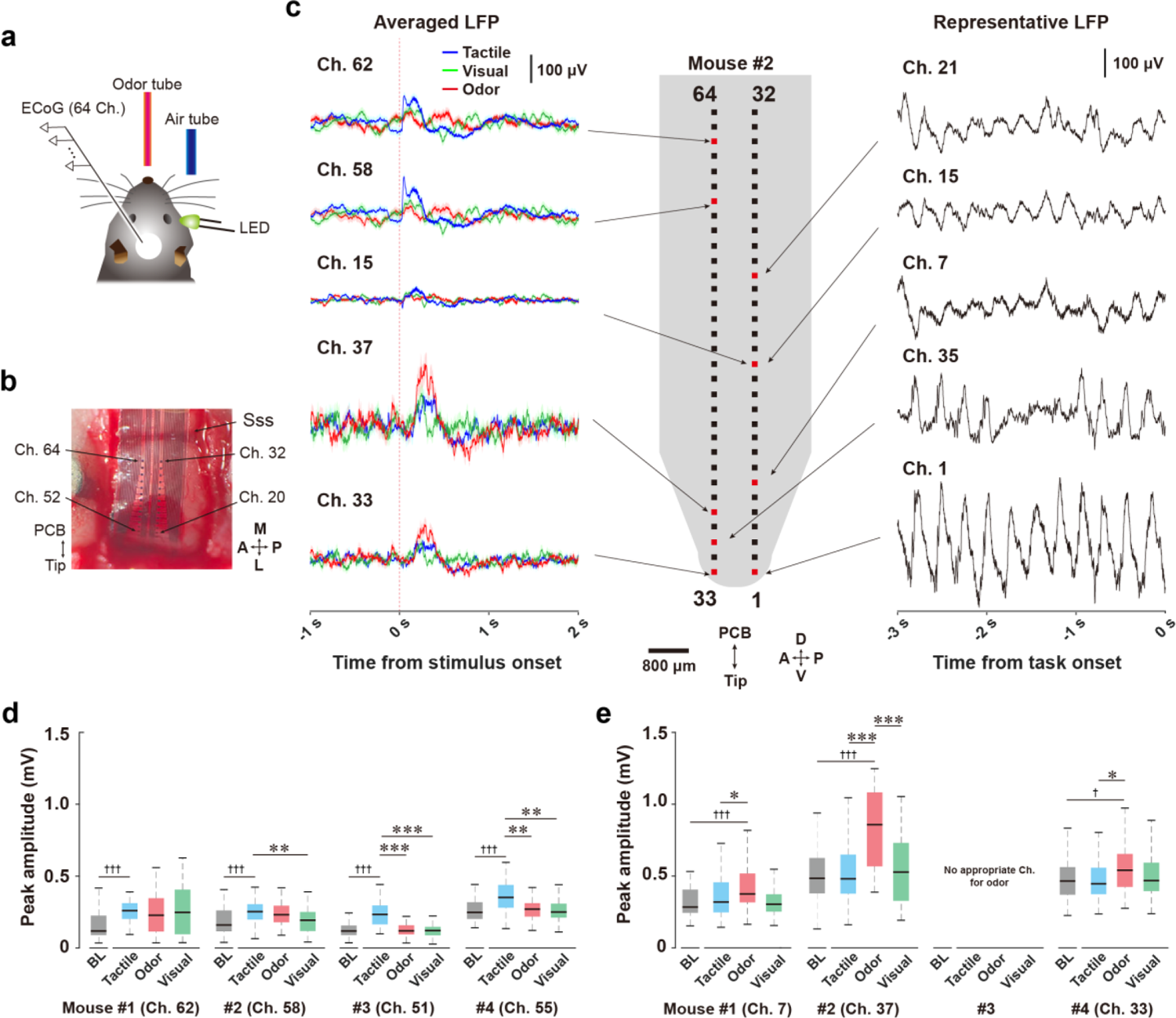
The μECoG electrode can monitor cortical activities in response to multiple sensory stimuli. (a) Schematic drawing of the sensory presentation task and recordings in a head-fixed mouse. (b) Dorsal view of the μECoG electrode array on the parietal cortex. M, medial; L, lateral; A, anterior; P, posterior; Ch., channel of electrode; Sss, superior sagittal sinus. (c) The 64-channel μECoG electrode array and LFPs at start of the task (representative LFP) or on task (averaged LFP) in a mouse. LFPs corresponding to the tactile, visual, and odor stimuli are indicated as blue, green, and red line, respectively. Shadows indicate SEM. D, dorsal; V, ventral; A, anterior; P, posterior. (d) Averaged peak LFP amplitudes recorded by the channels in which the maximal change was observed for the tactile stimulus. BL, baseline. (e) Averaged peak LFP amplitudes recorded by the channels in which the maximal change was observed for the odor stimulus. There are no channels showing significant differences between odor stimulus-evoked and baseline LFP amplitudes in the #3 mouse. The central lines and edges of the boxes indicate the median and quartiles, respectively, and whiskers extend to the maximum and minimum with outliers excluded. \**P* > 0.05, \*\*\**P* < 0.001 vs. each stimulus. †*P* > 0.05, ††† < 0.001 vs. baseline amplitude.

### Histological validation of μECoG sheet position

To examine whether the 23 μm flexible μECoG sheet appropriately reaches to the ventral parts of temporal cortex along the curved surface of mouse cerebral cortex, we histologically validated the position of the μECoG sheet by using adhesive transfer film to prevent the samples from separating [23] (Fig. 3a, see methods). Briefly, we prepared sections including brain, skull, and μECoG sheet and took microscopic images under conditions for bright field and DAPI (nuclear staining). The results showed that the μECoG sheet was situated along the surface of cerebral cortex (Fig. 3a, black arrows), and the tip of the μECoG sheet approached the piriform cortex, which is the most ventral part of the temporal cortex (Fig. 3a, red arrows). In addition, comparisons between ipsilateral and contralateral hemispheres showed no obvious damage in either the dorsal or ventral parts of the cerebral cortex that may have resulted from the surgical procedures required for device insertion (Fig. 3b-d).

### LFP Recordings in an awake mouse

First, we tested whether the μECoG electrode arrays, implanted onto the temporal cortex by our surgical methods, can monitor LFPs from a broad cortical area of an awake and head-fixed mouse, while also recording sensory stimulus-evoked potentials. Half of the electrodes on the μECoG sheet were inserted into the space between the temporal cortex and skull, and the remaining electrodes were situated on the parietal part of cortex (Fig. 4a). The location of the μECoG sheet in the temporal cortex was confirmed by a CT imaging after the experiment, and the tip of the sheet reached to the most ventral part of the temporal cortex (Fig. 4b), which is comparable with the histological assessment (Fig. 3a). Histological and CT assessments indicated that our μECoG electrodes were placed on parietal to temporal cortices, including the barrel and piriform cortices. Under this condition, we recorded LFPs with the μECoG electrode arrays (Fig. 4c) along with the dorsoventral axis of the cortex. Representative LFPs recorded by successive thirty-two electrodes are shown in Fig. 4d. The recording condition was reliably maintained for at least 3 hours under the acute experimental condition.

Next, we conducted a sensory cue-presentation task to record cortical potentials responding to sensory stimuli (Fig. 4e). In this experiment, we applied a tactile stimulus to the vibrissa in the form of mild air flow because the ECoG electrode arrays easily target the barrel cortex. The visual stimulus was used as a control stimulus as no electrodes were placed on the visual cortex, which is located in a more posterior region than the barrel cortex. Sensory stimuli were pseudorandomly presented to the mouse every ∼10–20 sec. The tactile stimulus induced strong tactile-evoked LFPs in the electrodes placed over the parietal region of the cortex, whereas the visual stimulus did not induce changes in the LFPs recorded from all channels (Fig. 4f). To examine the response characteristics to sensory stimuli on the same channel or between separated channels, we compared the amplitudes of maximal responses of another sensory stimulus and different electrodes to the tactile stimulus. We confirmed that channel 58 showed the maximal change in tactile stimulus-evoked LFP (Fig. 4g, Wilcoxon signed-rank test, *Z* = 6.009, *p* < 0.001, for comparison with baseline amplitudes), which was significantly higher than that of visual stimulus-evoked LFPs (Fig. 4g, Wilcoxon signed-rank test, *Z* = 6.087, *p* < 0.001). Moreover, we confirmed that the peak amplitudes of tactile stimulus-evoked LFP at channel 58 was significantly higher than that of channel 33 (Fig. 4h, Wilcoxon rank-sum test, *Z* = 5.112, *p* < 0.001), the most ventral electrode. To exclude the effects of currents originating far from the μECoG electrodes, we calculated the LFP profiles (see Method). The location of the largest sensory-evoked potential (Fig. 4f) and the largest LFP profile amplitude overlapped at channels surrounding channel 58, indicating that the surrounding region of channel 58 was highly active when the tactile stimulus was applied (Fig. 4i).

### LFP recordings in anesthetized mice

We tested whether the μECoG electrode array covering the parietal to temporal cortices can monitor cortical activity corresponding to multiple sensory stimuli. In this experiment, we conducted a sensory cue-presentation task that added an odor stimulus to the previous task that used tactile and visual stimuli (Fig. 5a). The neural recordings were carried out under urethane anesthesia because sniffing or chowing behaviors were frequently observed in mice not habituated to the novel odors of the task environments. As shown in Fig. 4a, approximately half of the electrode array on the μECoG sheet were inserted into the epidural space between the temporal cortex and skull, while the other electrodes were placed on the parietal cortex (Fig. 5b).

Representative LFP data were recorded by the μECoG electrode arrays from a wide area of cerebral cortex. Before the sensory cue-presentation task, we confirmed the presence of cortical LFPs from parietal to temporal cortices under anesthetized conditions (Fig. 5c, representative LFP). In the sensory cue-presentation task, tactile, odor, and visual stimuli were presented to the mouse pseudorandomly 50 times for each sensory modality (150 times total), every ∼10-20 sec. We observed clear tactile and odor stimulus-evoked LFPs from electrodes located on both parietal (Ch. 58 and 62) and temporal (Ch. 33 and 37) cortices, respectively (Fig. 5c, averaged LFP). Sensory stimulus-evoked LFPs were not observed from the electrode placed on the middle position of the recording areas along the dorsoventral axis (Ch. 15). We next measured the maximal amplitudes of the sensory stimulus-evoked LFP responses to the tactile, odor, and visual stimuli (50 trials each) in four mice. In channels placed on the parietal cortex, the tactile stimulus induced significantly higher evoked LFP amplitudes relative to the baseline amplitudes that were recorded just before the stimulus onset (Fig. 5d, Wilcoxon signed-rank test, *Z* = 4.609, *p* < 0.001 for Mouse #1, *Z* = 3.595, *p* < 0.001 for Mouse #2, *Z* = 6.042, *p* < 0.001 for Mouse #3, *Z* = 4.581, *p* < 0.001 for Mouse #4). The tactile stimulus did not result in higher evoked LFP amplitudes than visual and odor stimuli in the same channels in one mouse (Fig. 5d; Friedman test, *χ^2^* = 1.12, *p* = 0.571, for Mouse #1). In another mouse, the amplitudes evoked by the tactile stimulus were significantly higher than those evoked by the visual stimulus (Fig. 5d; one-way repeated ANOVA, *F*(2,147) = 4.86, *p* < 0.01, *post hoc* Tukey–Kramer test, tactile vs. odor, *p* = 0.805, tactile vs. visual, *p* < 0.01, odor vs. visual, *p* = 0.052, Mouse #2). In the remaining two mice, the tactile stimulus evoked higher amplitudes relative to the other stimuli (Fig. 5d; Friedman test, *χ^2^* = 39.04, *p* < 0.001, *post hoc* Tukey–Kramer test, tactile vs. odor, *p* < 0.001, tactile vs. visual, *p* < 0.001, odor vs. visual, *p* = 0.916 for Mouse #3; *χ^2^* = 14.68, *p* < 0.001, *post hoc* Tukey–Kramer test, tactile vs. odor, *p* < 0.01, tactile vs. visual, *p* < 0.01, odor vs. visual, *p* = 0.916 for Mouse #4). The amplitudes of odor stimulus-evoked LFP recordings from channels located on the ventral parts of the temporal cortex were significantly higher than the baseline amplitude in three mice, but not in one mouse (Mouse #3) (Fig. 5e; Wilcoxon signed-rank test, *Z* = 3.464, *p* < 0.001 for Mouse #1, *Z* = 5.146, *p* < 0.001 for Mouse #2; paired Student’s *t*-test, *t*_(49)_ = 2.471, *p* < 0.05 for Mouse #4). In the three mice, the odor-evoked LFP amplitudes tended to be higher than other stimuli-evoked LFP amplitudes (Fig. 5e). Two of the mice showed that the odor stimulus-evoked amplitudes were larger than either the visual or tactile-evoked LFP amplitudes (Fig. 5e; Friedman test, *χ^2^* = 8.32, *p* < 0.05, *post hoc* Tukey–Kramer test, tactile vs. odor, *p* = 0.112, tactile vs. visual, *p* = 0.703, odor vs. visual, *p* < 0.05 for Mouse #1; *χ^2^* = 6.28, *p* < 0.05, *post hoc* Tukey–Kramer test, tactile vs. odor, *p* < 0.05, tactile vs. visual, *p* = 0.514, odor vs. visual, *p* = 0.341 for Mouse #4). On the other hand, the remaining mouse exhibited odor stimulus-evoked amplitudes that were larger than either the visual or tactile-evoked LFP amplitudes (Fig. 5e; Friedman test, *χ^2^* = 20.32, *p* < 0.001, *post hoc* Tukey–Kramer test, tactile vs. odor, *p* < 0.001, tactile vs. visual, *p* = 0.978, odor vs. visual, *p* < 0.001 for Mouse #2).

## Discussions

EEG and ECoG recordings in humans and non-human primates are used to record neural activity from a wide area of cerebral cortex, including the temporal lobe, with high temporal and spatial resolution [24, 25]. However, there are no techniques to record from the temporal cortex using these techniques in rodents. Such a techniques would provide a powerful tool for neuroscience. Here, we describe the development of a novel μECoG electrode array, including surgical techniques for implanting the device into the mouse temporal cortex. We also confirm that large-scale LFP recordings can be obtained from the novel μECoG electrode array. We show that array thickness is a key factor for the usefulness of parylene-based electrodes. Appropriate stiffness of the μECoG sheet-shaped electrode allowed its insertion into the space between the skull and brain surface, while minimizing the invasiveness of the arrays on the mouse cerebral cortex. In addition, the device could reliably assess sensory stimulus-evoked LFPs from the multiple recording sites covering the somatosensory and olfactory cortex both in awake and anesthetized mice. These techniques are suitable for understanding mechanisms relevant to cognitive functions and pathological conditions in which inter-regional network activities are involved.

Tissue penetrating neural electrode arrays are the primary method used for in vivo neuronal recordings in rats and mice. However, damages to brain tissue or dura mater cannot be avoided, and glial scar formation can occur around electrode arrays, leading to quality degradation of neural signals [26]. By contrast, sheet-shaped ECoG electrode arrays are less invasive than penetrating electrode arrays. Several groups have developed insertable devices that implant onto the brain surface, beneath the skull [17, 18, 27, 28]. Although these techniques enabled neural recordings from a relatively wide area of the cerebral cortex without penetrating the brain and with little surgery-related tissue trauma, the recording areas were limited to the dorsal parts of cerebral cortex in rodents. Our μECoG sheet overcomes this limitation by expanding the area of cortical recordings to other areas. Using CT imaging and histological methods, we confirmed that placement of the μECoG sheet-shaped array covered the area from the somatosensory cortex to the piriform cortex [29] without structural damage to the cortical surface. In addition, odor stimulus-evoked LFPs were observed from electrodes placed on the tip of the array. These results strongly demonstrate that our μECoG can record from a wider area of the mouse cerebral cortex than existing ECoG methods., This novel μECoG method may shed light on neural network interactions between the parietal and temporal cortex in mice.

In this study, we characterized the performance of our μECoG device under awake and anesthetized conditions. The next step would be to assess the usefulness of the device under chronic conditions. The μECoG device was fabricated using parylene, which is a biocompatible material. Previous studies have indicated that parylene-based μECoG devices can maintain electric impedance and the quality of chronic recordings for up to two weeks [30]. Additionally, our surgical methods enabled implantation of the electrode array without structural damage to the dura matter or the layered tissue structures of the cerebral cortex (Fig. 3b-d), despite the wide-area craniotomy needed for electrode placement (Fig. 1g, 4a, and 5b). Our results are consistent with those of a previous study describing a methodology for implanting an μECoG device into the space between the brain surface and skull [18]. Together, the characteristics of our device and surgical methods suggest that they can be used in chronic applications. On the other hand, chronically implanted μECoG devices can cause micro-hematomas and result in opaque connective tissue, which can affect normal cortical activity [31]. Further studies should investigate whether the signal quality is maintained several weeks after implanting the μECoG device using our novel methodology.

Thickness is correlated with stiffness and is one of the key factors for the development of thin-film biomedical devices that adhere appropriately to the complex curvature of the skin, heart, and brain surface [6, 18, 32]. Stiffness and adhesion are trade-offs that need be considered in thin film development; for instance, a device with high adhesion properties is not appropriate for insertion into narrow spaces such as the epidural space. Indeed, the insertable µECoG was relatively thicker than the ECoG devices reported previously [6], and therefore, it was possible to move the electrodes to the target area in the narrow space between skull and cortex. Furthermore, the recording data obtained by the ECoG device were comparable to those of devices developed previously [16, 33]. This indicates that optimization of thickness and stiffness is important for the development of adequate devices.

Brain-machine interfaces (BMIs) and brain-computer interfaces (BCIs) allow interactions between the brain and external devices [34]. These technologies are expected to restore lost function in patients with neurological impairments. The ECoG has the advantage of being less invasive than penetrating types of electrodes. Another advantage is to be more informative than EEG because of the higher spatial resolution [35]. Previous studies reported that the ECoG signals related to sensory and motor information processing are sufficient to decode the movement trajectory of one- to three-dimensional space, and to manipulate a cursor on a computer screen and a robotic arm [36–38]. In addition, readout of higher-order cognitive brain signals is expected to advance cognitive BMI technology [39]. Applying the µECoG to rodents will boost development of the BMIs/BCIs technology because it is easier to apply to rodents than to humans and non-human primates. Given that the temporal cortex in rodents, including the insular and piriform cortices, is involved in decision making processes based on sensory information [12, 40], enlarged recording techniques covering the parietal to temporal parts of cortices in mouse might have potential for the development and implementation of cognitive BMIs.

ECoG and μECoG techniques can be employed in humans, non-human primates, and rodents using a similar methodology [6, 24, 25], thus providing physiological data that can be compared across species. However, the measurement area of current μECoG techniques in mice is limited to the relatively small cortical surface area, and currently does not include the temporal and occipital cortex, which are easily addressed in humans and non-human primates. The μECoG device developed here is insertable into the most ventral parts of the epidural space and makes it possible to record cortical activity from the dorsal to ventral cortices of mice, which has previously been difficult to achieve. Our approach extends the possibility of comparing features of ECoG recordings between mouse and human and non-human primates. Therefore, our μECoG device and methodologies could promote translational research by applying them to rodent models of various pathologies. Possibilities include the combination of μECoG recordings with optogenetics to investigate the causes of diseases. In the future, the integration of these genetical tools and large-scale μECoG recordings may help to further elucidate the mechanisms underpinning neuropsychiatric disorders. Indeed, neuropsychiatric disorders of the brain have been traditionally understood as loss or disruption of brain function due to changes in cell function, neurotransmitter systems, or perturbed neuronal connections. Previous reports have reported changes in alpha and theta oscillation in a wide area of the cortex in Alzheimer’s disease patients [41, 42]. Additionally, EEG studies have reported impairments in synchronization between gamma and neural oscillations across various cortical region [43–45]. Although several technologies have been developed to record neural activities in a wide area of the brain, μECoG provides better spatial resolution and signal to noise ratio than EEG and functional magnetic resonance imaging. On the other hand, optogenetics have been used to manipulate selectively cell types and neural circuits. Thus, a combination of μECoG and optogenetics could find out principles of diseases by monitoring and by adding manipulation to impaired neural activities of wide area of cortex in order to promote translational research and/or restore neuropsychiatric diseases.

## Conclusion

In this study, we developed a 64-channel μECoG device covering the parietal to temporal cortex in mice. The novel surgical procedures enable device placement on the piriform cortex, which is the most ventral part of the mouse cerebral cortex. The device recorded stimulus-evoked potential across multiple sensory areas of cerebral cortex along the dorsal-ventral axis. This system will provide opportunities to investigate physiological functions from wider areas of the mouse cerebral cortex than those of existing ECoG techniques.

## Abbreviations

mECoG: micro (m)-electrocorticography

CT: computed tomography

EEG: electroencephalography

LFP: local field potential

PCB: printed circuit board

DAPI: 4’,6-diamidino-2-phenylindole

## Declarations

### Ethics approval and consent to participate

All animal procedures were approved by the Animal Care and Use Committee of Dokkyo Medical University (approval number, 1339) and carried out in accordance with the guidelines of the National Institutes of Health.

## Consent for publication

Not applicable.

## Availability of data and materials

All data used in this paper are available upon reasonable request to the corresponding author.

## Author contributions

S.S., H.S., and N.O. conceived the project. O.F. and M.O. evaluated the characteristics of the μECoG device. S.S. and Y.S. performed the animal experiments. S.S. analyzed the data. M.I., S.N., and T.H. performed the histological experiments. R.K. and S.T. performed the fabrications and its measurements for the μECoG device. H.M. and K.M. supported the developments for behavioral apparatus and systems. F.S. and K.H. scanned CT imaging. S.S., R.K., H.S., and N.O. wrote the manuscript.

## Acknowledgments

We thank the Research Center for Laboratory Animals, Dokkyo Medical University for animal maintenance, the Center for Research Collaboration and Support, Dokkyo Medical University for allowing to use the BZ-X700 fluorescence microscope, and S. Fujiki and all members of Division for Memory and Cognitive Function, Dokkyo Medical University for valuable discussions.

