## Additional file for "A novel micro-ECoG recording method for recording multisensory neural activity from the parietal to temporal cortices in mice"

**This file includes:**

Additional Method

Fig. S1

### Additional Method

##### Channel crosstalk

The crosstalk between electrode channels was investigated using an equivalent circuit model as shown in Fig. S1a. This circuit models two adjacent recording channels, where $V_{\mathrm{i1}}$ and $V_{\mathrm{i2}}$ are input voltages of each channel, $V_{\mathrm{o1}}$ and $V_{\mathrm{o2}}$ are output voltages of each channel, $Z_{\mathrm{e}}$ is electrode-electrolyte impedance, $Z_{\mathrm{c}}$ is coupling capacitance impedance, $Z_{\mathrm{p}}$ is parasitic capacitance impedance to ground, and $Z_{\mathrm{A}}$ is input impedance of amplifier, respectively. The cross talk is evaluated by the ratio of the signal on the input voltage $V_{\mathrm{i1}}$ to the output voltage $V_{\mathrm{o2}}$ of the adjacent channel when $V_{\mathrm{i2}} = 0$. In an ideal circuit, since $Z_{\mathrm{p}}$ and $Z_{\mathrm{A}}$ can be removed as infinite, the crosstalk can be easily evaluated as expressed in the following equation:

$$\frac{V_{\mathrm{o2}}}{V_{\mathrm{i1}}} = \frac{Z_{\mathrm{e}}}{2Z_{\mathrm{e}} + Z_{\mathrm{c}}}$$

Next, we consider the value of $Z_{\mathrm{c}}$ from the design of the μECoG sheet. The coupling capacitance $C_{\mathrm{c}}$ can be expressed:

$$C_{\mathrm{c}} = \varepsilon_{\mathrm{p}} \frac{tL}{s}$$

where $\varepsilon_{\mathrm{p}}$ is the permittivity of dielectric (parylene C), *t* and *L* are the thickness (250 nm) and length (15 mm) of metal wiring, and *s* is spacing (20 μm) between the metal wiring layer, respectively. Based on the device parameters, the $C_{\mathrm{c}}$ is calculated to be 5.1 fF, which corresponds to the impedance $Z_{\mathrm{c}}$ of 31 GΩ at 1 kHz. The average $Z_{\mathrm{e}}$ in the μECoG sheet as shown in Fig. 2d was 0.5 MΩ. Therefore, the crosstalk $V_{\mathrm{o2}}/V_{\mathrm{i1}}$ is calculated to be $1.61 \times 10^{-5}$ (−48 dB). There is almost no influence of channel crosstalk in the developed μECoG sheet. Additionally, the effect of spacing between electrode channels on the calculated crosstalk for $Z_{\mathrm{e}}$ of 0.5 and 5 MΩ was calculated as shown in Fig. S1b. The crosstalk increases with decreasing spacing and impedance. Therefore, the μECoG sheet with low impedance electrode and wide spacing are more effective for ECoG measurement.

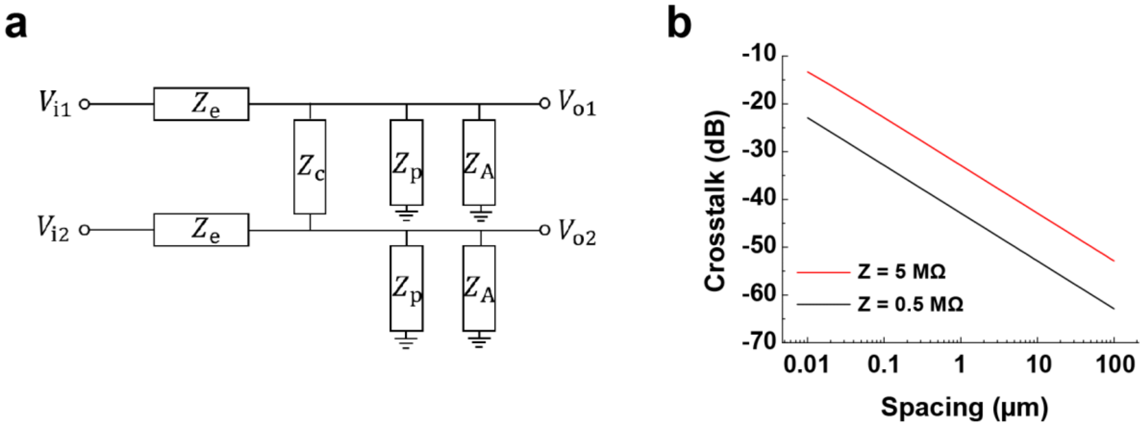


**Fig. S1. Channel crosstalk.** (a) Equivalent circuit model representation of two adjacent recording channels with electrode-electrolyte impedance $Z_e$, coupling capacitance impedance $Z_c$, parasitic capacitance impedance to ground $Z_p$, and input impedance of amplifier $Z_A$. (b) Crosstalk as a function of spacing between two electrode channels at different electrode impedance $Z_e$ of 0.5 and 5 MΩ with thickness *t* of 0.25 mm, length *L* of 15 mm of metal wiring layer, and spacing *s* of 20 μm.
